# Epigallocatechin Gallate Modulates Microglia Phenotype to Suppress Pro-Inflammatory Signalling Cues and Inhibit Phagocytosis

**DOI:** 10.1101/2023.05.24.542060

**Authors:** Philip Regan, Katriona L Hole, Julia Sero, Robert J Williams

**Affiliations:** Department of Life Sciences, University of Bath, Bath, UK

**Keywords:** Microglia, Alzheimer’s Disease, Flavonoids, Phagocytosis, Amyloid beta

## Abstract

Microglia are crucial players in the pathogenesis of late onset Alzheimer’s Disease (AD), with evidence for both deleterious and beneficial effects. Identifying interventions to modulate microglial responsiveness, to promote Amyloid β (Aβ) clearance, disrupt plaque formation or to dampen excessive inflammation has therapeutic potential. Bioavailable flavonoids such as the flavan 3-ols are of interest due to their antioxidant, metal chelating, signalling and anti-inflammatory potential. Primary microglia were treated with a series of structurally related flavanol 3-ols to assess effects on phagocytosis, cytokine release and transcriptional responses by RNA sequencing. Data indicated that the extent of hydroxylation and the presence of the galloyl moiety were strong determinants of flavan 3-ol activity. Epigallocatechin gallate (EGCG) was the most effective flavan-3-ol tested and strongly inhibited phagocytosis of Aβ independent of any metal chelating properties suggesting a more direct modulation of microglia responsiveness. EGCG was broadly anti-inflammatory, reducing cytokine release and downregulating transcription particularly of components of the microglia extracellular matrix such as MMP3 and SerpinB2. Collectively this brings new insight into the actions of flavonoids on microglial responsiveness with potential implications for the therapeutic use of EGCG and structurally related flavanol-3-ols in AD.

## Introduction

Alzheimer’s Disease (AD) is characterized by a long prodromal phase, during which time amyloid β-peptide (Aβ) accumulates, aggregates and ultimately deposits as plaques within the brain. This process triggers the emergence of tau pathology and microglial driven inflammatory responses. Microglia are recruited to Aβ plaques where they acquire and express a disease-associated microglia (DAM) transcriptional signature [1]. Plaque-associated microglia potentially sequester Aβ aggregates and restrain Aβ, thereby delaying disease progression, although microglia might equally be required for parenchymal plaque formation [2,3]. Excessive microglial activation appears to exacerbate neuronal damage particularly during later stages of AD [3–5] and genome wide association studies strongly implicate microglia in AD risk [6], so overall the role of the innate immune system is complex. Nonetheless, identifying strategies to modulate microglial responsiveness, either to promote Aβ clearance, disrupt plaque formation or to dampen excessive inflammation is of therapeutic interest, although the correct timing of any neuroimmune intervention will be critical to any successful outcome.

The underlying reasons why Aβ levels increase and aggregate in sporadic AD is not clear but could be linked to a progressively pro-oxidant environment in the aged brain and/or the presence of redox sensitive metals. This hypothesis has generated considerable interest in the use of flavonoids and other dietary polyphenols in AD, due to their antioxidant, metal scavenging and secretase modulating activity [7–10]. Some flavonoids also directly disrupt Aβ aggregation by favoring the formation of unstructured off-target oligomers [11]. However, despite possessing impressive efficacy in vitro, very few flavonoids have progressed beyond early phase preclinical development due to considerable challenges around absorption, distribution, metabolism, and excretion (ADME)[12]. The group of flavonoids with the best combination of bioavailability and activity are the flavan-3-ols which possess a common 2-phenyl-3,4-dihydro-2H-chromen-3-ol skeleton, but with varying numbers of hydroxyl groups and galloyl moieties that together characterise the major individual flavan-3-ol monomers. Epigallocatechin gallate (EGCG) is a bioactive flavonoid belonging to this subfamily which has progressed to phase 2/3 clinical trials for AD and Down’s syndrome [13] based on a multi-modal activity profile including anti-Aβ, anti-inflammatory actions and regulation of the unfolded protein response [14]. These effects may extend to direct modulation of microglia function, which could therefore, underly reported protective effects of EGCG and other flavan-3-ols for the brain. On this basis we undertook structure-activity studies to establish whether flavan-3-ols directly modulate the phagocytosis function of primary mouse microglia, and whether structural characteristics of flavan-3-ols differentially affect microglia function.

Microglia were treated with a structurally related series of flavan-3-ols, and data indicated that hydroxylation and the presence of the galloyl moiety were strong determinants of their microglia modifying properties. EGCG was the most effective flavan-3-ol tested and inhibitory actions on phagocytosis activity were independent of metal chelating properties, suggestive of a more direct modulation of microglia responsiveness. EGCG was broadly anti-inflammatory, reducing cytokine release and downregulating transcription particularly of components of the microglia extracellular matrix. Collectively this brings new insight into the actions of EGCG on microglial responsiveness with potential implications for the therapeutic use of EGCG and structurally related flavanol-3-ols in AD.

## Materials and Methods

### Animals

All procedures involving animals were carried out in accordance with the UK Animals Scientific Procedures Act, 1986. 1–3-day old male CD1 mice were used to prepare primary cultured glia. All animal procedures were given ethical approval by the Animal Welfare and Ethical Review Body at the University of Bath.

### Flavonoids

All flavonoids described in this study were purchased from Extrasynthese (France) and solubilised in dimethyl sulfoxide (DMSO) to prepare a 10mM stock solution that was then used to derive final working concentrations.

### Mixed Glia Primary Culture

Following cervical dislocation, brains were removed from mouse pups and placed into ice-cold HBSS (#14175095, Fisher Scientific) and whole cortex was dissected under sterile conditions. Tissue was dissociated through incubation with 0.25% Trypsin (#15090046, Fisher Scientific) at 37°C for 15 min and then 0.2mg/ml DNase I (#DN25-100MG, Merck) at room temperature for 5 min, followed by repeated trituration using fire-polished Pasteur pipettes. Cells were resuspended in Glia Feed Media (DMEM/F12 without HEPES or phenol red (#11580546, Fisher Scientific), supplemented with 10% FBS (#10500064, Fisher Scientific), 1% Penicillin/Streptomycin (#15140122, Fisher Scientific) and 1x Glutamax (#11574466, Fisher Scientific), then passed through a 70µm cell strainer (# 15346248, Fisher Scientific), plated onto 60mm dishes pre-coated with poly-L-ornithine (10ug/ml; #P4957-50ML, Merck), and stored in a 37°C incubator with 5%CO_2_.

A full media change was completed at DIV 1, 2/3 feed at DIV 5 and 1/2 feed every subsequent 3 days with Glia Feed Media. Following establishment of an astrocytic monolayer at around DIV 12, media was supplemented with 10ng/ml GM-CSF (#315-03, Peprotech) to promote microglia proliferation.

### Isolation of Microglia

Microglia were isolated from mixed glia cultures at DIV20-30, using shaking to detach surface microglia (for 96-well plate assays) and/or trypsinisation to remove the astrocytic layer (for RNA extraction and conditioned media assay). To detach surface microglia, mixed glia cultures were shaken for 5 min at 200RPM followed by vigorous tapping. Collected microglia were re-plated into black-walled 96-well plates (#M0562-32EA, Greiner) coated with 50ug/ml poly-D-lysine (#P7280-5MG, Merck) at 18,000 cells/well in a 1:1 mix of conditioned and fresh Glia Feed Media. For astrocyte detachment, cells were washed with HBSS and then incubated with 0.083% Trypsin-EDTA (#25200056, Fisher Scientific) at 37°C for 30-45 min. Remaining microglia cells were washed and then re-fed with a 1:1 mix of conditioned and fresh Glia Feed Media.

24 h after isolation, microglia were washed with HBSS and media was replaced with Serum Free Microglia Media (DMEM/F12 without HEPES or phenol red, supplemented with 1% Penicillin/Streptomycin, 1x Glutamax, 1x Insulin-Transferrin-Selenium (#12097549, Fisher Scientific), 0.1% BSA (#A8806-1G, Merck), 100ng/ml IL-34 (#200-34, Peprotech), 2ng/ml TGFβ2 (#100-35B, Peprotech), 1ng/ml GM-CSF and 300ng/ml cholesterol (#700000P-100MG, Merck). Microglia were left in Serum Free Microglia Media for 24 h prior to addition of treatments for experimental assays.

### Phagocytosis Assays

Lyophilised HiLyte Fluor488 labelled amyloid-β_1-42_ peptide (Aβ_42_) (#AS-60479-01, Cambridge Biosciences) was resuspended to 1mg/ml in 100% 1,1,1,3,3,3-Hexafluoro-2-propanol (HFIP), and the mixture was vortexed occasionally during 1 h to ensure proper solvation. The solution was then aliquoted into separate vials and HFIP was allowed to evaporate in the fume hood overnight, after which vials were sealed and stored at −20°C protected from light. Before use, the peptide film was brought to room temperature and resuspended in 100% anhydrous DMSO to make a 1mM solution. The peptide solution was left for 15 min, with occasion vortexing to ensure dissolution. D-PBS was added to prepare a 100µM working solution that was then left to incubate at room temperature for 2 h.

pHrodo Green Zymosan Bioparticles (#P35365, Fisher Scientific) were resuspended to 0.5mg/ml in Live Cell Imaging Solution (#12363603, Fisher Scientific), followed by alternate trituration and vortexing for 10 min, immediately prior to use.

On the day of the assay, pre-treated microglia in 96-well plates were switched to pre-warmed Live Cell Imaging Solution supplemented with 0.9% D-(+)-Glucose, CellMask Deep Red (1:1000; #C10046, Fisher Scientific) and NucBlue Live ReadyProbes Reagent (#R37605, Fisher Scientific), and left at 37°C for 20 min to achieve live-cell staining. Following staining, microglia were switched to pre-warmed Live Cell Imaging Solution supplemented with 0.9% D-(+)-Glucose, and HiLyte Fluor488 Aβ_42_ (final concentration 500nM) or pHrodo Green Zymosan bioparticles (final concentration 33µg/ml) were added to relevant wells. Plates were placed into an IN Cell Analyzer 2000 (GE Healthcare) at 37°C for live-cell imaging of uptake every 30 min for a total of 2 h.

### Immunocytochemistry

Microglia in 96-well plates were washed twice with D-PBS, before fixation with 4% PFA and 4% sucrose in D-PBS for 20 min. After washing, cells were permeabilised with 0.15% Triton X-100 for 15 min, and then incubated with blocking solution (2.5% Normal Goat Serum and 1% BSA in D-PBS) for 30 min. Primary antibodies were added to blocking solution and incubated overnight at 4°C after which cells were washed and incubated with relevant fluorescently labelled goat-derived secondary antibodies (1:300) in blocking solution for 1 h at room temperature. Finally, cells were incubated with a nuclear counterstain (NucBlue Live ReadyProbes Reagent) and images were acquired using the IN Cell Analyzer 2000. The following primary antibodies were used: rat anti-CD11B (1:100; #MCA711G Bio-Rad), rabbit anti-IBA-1 (1:100; #AB178847 Abcam), mouse anti-NFκB p65 (F-6) (1:100; #sc-8008 Santa-Cruz).

### Nitrite release assay and inflammatory stimuli

Nitrite levels in microglia conditioned media were assessed using the Promega Griess Reagent system (#G2930) according to the manufacturer’s instructions. 50µl/well of conditioned culture media was taken in duplicate from microglia treated for 24h in a 96-well plate. Samples were mixed with sulfanilamide and N-1-naphthylethylenediamine dihydrochloride (NED) solutions in the assay plate and absorbance read on a plate reader. LPS (0.1µg/ml) (O111:B4 #L4391-1MG, Sigma-Aldrich) and 20ng/ml Interferon-γ (IFN-γ; 315-05, Peprotech) were used as inflammatory stimuli. The selective inhibitor of nitric oxide synthase, L-NIL hydrochloride (30µM), was used as a negative control.

### Image Acquisition and Analysis

Automated image capture was achieved using the IN Cell Analyzer 2000, using a 40x objective and 5-6 regions of interest per well and custom acquisition settings per fluorophore. Following capture, images were pre-processed using ImageJ and CellProfiler software to correct for flatfield background illumination artifacts. A custom CellProfiler pipeline was optimised to provide primary and secondary object identification in each image, thereby identifying individual nuclei and their associated cell bodies from nuclear and cytoplasmic/membrane specific stains, respectively. Quantitative measures of stain intensity in nuclear/cytoplasmic cell compartments, and of cell features (e.g., morphology) on a per cell basis were output to .CSV files through the CellProfiler pipeline. Finally, a custom *R* script was used for data quality control, analysis, and visualisation.

### RNA extraction and Sequencing

Microglia RNA was extracted using the RNeasy mini kit (Qiagen) following the manufacturer’s instructions, including on column DNase digestion. Purified RNA samples were sent to Novogene UK for quality control and subsequent strand-specific mRNA quantification using the NovaSeq PE150 platform (40M reads per sample). Briefly, raw reads were filtered for adapter contamination and low-quality reads, and sequence alignment was performed with HISAT2 to map clean reads to the reference mouse genome (GRCm38.p6). Raw counts of sequencing reads were used for subsequent differential expression analysis. Raw and processed data files are uploaded to the GEO server and available to access using the GEO accession number GSE208144.

### RNA-sequencing analysis

Read counts were input into a custom *R* workflow, in which data was initially filtered to remove low count genes (< 10 reads) and non-protein-coding genes, after which the DESeq2 R package (v1.30.1) was used to analyse differential expression across treatment groups. Differential expression analysis was performed on the linear modelled dataset with the experimental design matrix representing the difference between ‘EGCG and ‘Vehicle’ treatments. Wald test was used for hypothesis testing when analysing log2 fold changes for each gene between conditions, and the Benjamini-Hochberg method used to correct for multiple testing, with adjusted p-values (FDR)<0.1 considered significant.

A gene significance score, or π-value, was calculated for each gene by multiplying the log2 fold change by the -log10 *P*-value [15] which was then used for gene ranking and applied to gene set enrichment analysis for Gene Ontology (GO) with GSEA using the R package clusterProfiler (v3.18.1) The GO annotation (Biological Process (BP), Molecular Function (MF) and Cell Compartment (CC)) was mapped to gene sets consisting of 15-500 genes obtained from Molecular Signatures Database (MSigDB v7.1) and enrichment analysis was tested using 1000 permutations and an FDR Q-value of 0.1 used to identify significantly enriched gene sets in EGCG vs Vehicle treatment conditions. A gene enrichment ratio was calculated as the proportion of DEGs within a given gene set and only GO terms with ratios > 0.15 were visualised.

### Conditioned Media Proteome Profiler

Microglia in Serum Free Microglia Media were treated for 24 h with 1µM EGCG, 20ng/ml IL-3 (#200-03, Peprotech), or 20ng/ml IL-4 (#200-04, Peprotech) and media was subsequently collected into pre-chilled Eppendorf tubes, centrifuged at 4°C for 10 min to remove cell debris and media was stored at −80°C until use. Media collected from 3 biological replicates were pooled and a total of 500µL was added to a Mouse XL Proteome Profiler Cytokine Array (#ARY028, R&D Systems) according to the manufacturer’s instructions. Following background subtraction, densitometry of individual blots was measured using the AnalyzeDotBlot macro script in ImageJ and subsequent data analysis was performed using a custom R script.

### Statistical Design and Analysis

A two-way ANOVA or Mixed effects model (not assuming sphericity) was used to test for effects of treatment and/or time on microglia phagocytosis assays, followed by Tukey’s or Šídák’s multiple comparisons test to determine statistically significant effects between specific groups. GraphPad Prism v9 was used for data analysis including non-linear regression curve fitting for dose-response analysis.

## Results

### Structure-inhibitory activity of flavan-3-ols for modulation of microglia phagocytosis

Microglia isolated from the mouse brain and maintained in culture expressed the microglia-specific proteins C11b and Iba1 as shown using immunocytochemical approaches (**Fig. 1A**). These cells also exhibited functional characteristics expected of microglia as effectors of the innate immune response in the central nervous system (CNS). Consistent with their pivotal role in inflammatory signalling, robust nuclear translocation of the transcription factor, NFκB, was demonstrated in microglia following exposure to pro-inflammatory stimuli (LPS + IFNL; unpaired t-test: t(8) = 6.3, P=0.0002) (**Fig. 1B**). Similarly, release of pro-inflammatory nitric oxide (NO) from microglia was strongly induced by LPS + IFNL treatment, in a NO synthase dependent manner (one-way ANOVA F(2,19) = 50.2, Vehicle vs LPS + IFNL P < 0.0001) (**Fig. 1C**). Cultured primary microglia exposed to fluor 488-labelled Aβ_42_ actively internalised Aβ through phagocytosis, consistent with their proposed role in Aβ clearance in the brain (**Fig. 1D**). Collectively, this validated our model as a suitable platform for exploring the bioactivity of flavonoids against key aspects of microglia function relevant to AD.

**Figure 1.**
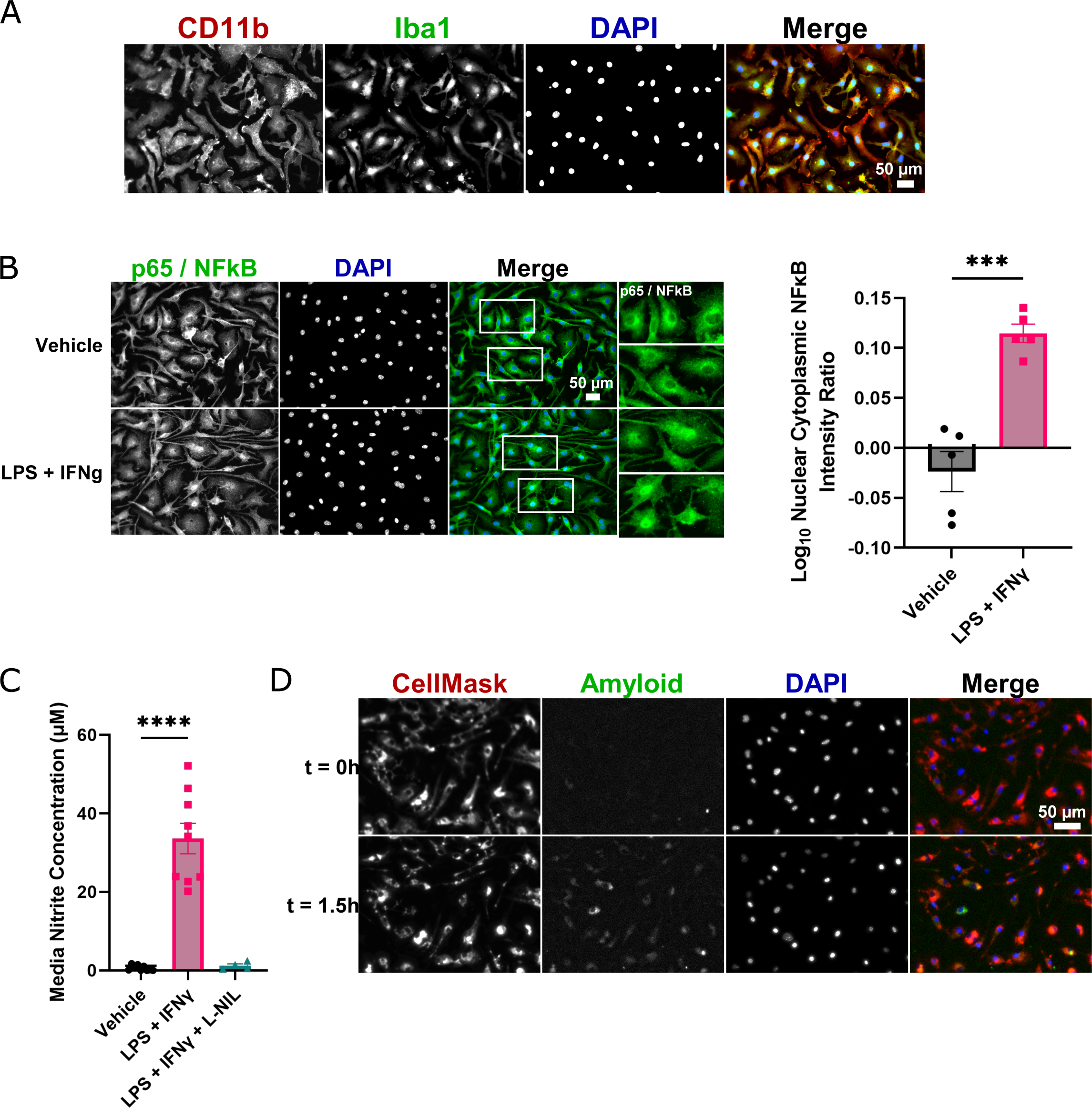
Phenotypic characterisation of primary microglia. (A) Immunofluorescence images of primary microglia stained with antibodies to the microglia markers CD11b and Iba1. (B) Immunofluorescence images of primary microglia stained with NFκB p65 antibody following 24h exposure to 0.1µg/ml LPS and 20ng/ml IFN-L. White boxes indicate inset images. Quantification of the log ratio of nuclear:cytoplasmic NFκB for each treatment group is shown in the right panel. Datapoints and error bars indicate mean ± S.E.M. *** < 0.005, (C) Quantified media nitrite levels following microglia 24h exposure to 0.1µg/ml LPS and 20ng/ml IFN-L. Co-treatment with L-NIL (30µM) in a subset of wells was used as a negative control treatment group. Datapoints and error bars indicate mean ± S.E.M. **** < 0.0001, (D) Representative images of primary microglia live-stained with a nuclear marker (DAPI; blue) and a cell membrane marker (CellMask; magenta), showing phagocytosis of 500nM HiLyte Fluor488 labelled amyloid-beta_1-42_ peptide (green) 0h and 1.5h after exposure.

We first tested whether flavan-3-ols modulate the phagocytosis function of microglia, and whether structural characteristics of flavan-3-ols differentially affect microglia function. Microglia in serum free media were treated for 24 h with 10µM of the indicated flavan-3-ols (**Fig. 2A**) and then exposed to 500nM fluor 488-labelled Aβ_42_ in their presence. Uptake of Aβ_42_ was quantified periodically by measuring the average 488-fluorescence intensity of individual microglia (i.e., the amount of internalised Aβ_42_ per microglia) and the percentage of microglia exhibiting above-background 488-fluorescence (i.e., the proportion of actively phagocytosing microglia) (**Fig. 2B**). Our data show that, relative to vehicle treatment, the average 488-fluorescence intensity was reduced in microglia in a flavan-3-ol structure-dependent manner (two-way ANOVA effect of treatment: F(5,12) = 11.17, P = 0.0004) (**Fig. 2C**). EGCG was the most efficacious inhibitor of microglia phagocytosis of Aβ_42_, with a significant reduction in the amount of Aβ_42_ internalisation from 1h onwards (Tukey’s multiple comparison Vehicle vs EGCG: 1h P = 0.0023, 1.5h P = 0.001, 2h P < 0.0001) (**Fig. 2E**). The proportion of phagocytosing microglia was also significantly reduced by EGCG compared to other flavan-3-ols (two-way ANOVA effect of treatment: F(5,12) = 8.09, P = 0.0015; Tukey’s multiple comparison Vehicle vs EGCG: 1h P = 0.001, 1.5h P < 0.0001, 2h P = 0.005) (**Fig 2D, F**). These data indicate that hydroxylation and the presence of the galloyl moiety are strong determinants of the microglia modifying properties of flavan-3-ols.

**Figure 2.**
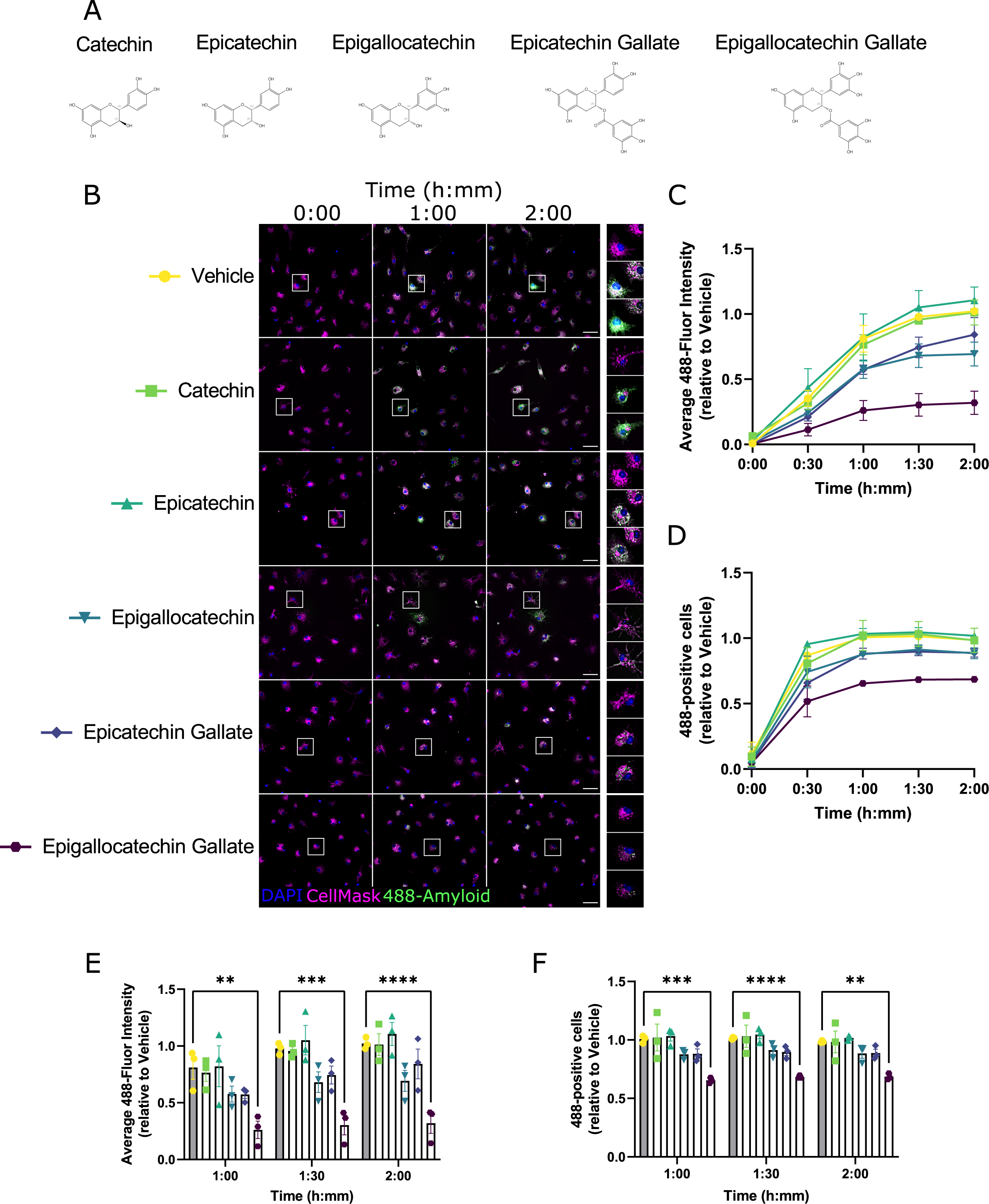
Structure-response screen for flavan-3-ol modulation of microglia phagocytosis. (A) Chemical structures of selected flavan-3-ols used in this study. (B) Representative immunofluorescence images of primary microglia live-stained with a nuclear marker (DAPI; blue) and a cell membrane marker (CellMask; magenta), showing phagocytosis of HiLyte Fluor488 labelled amyloid-beta_1-42_ peptide (green) at different timepoints. Individual rows represent individual treatment groups. White boxes indicate inset images on the right panel. Scale bar = 50µm. (C) Quantified time course data of average cellular 488-fluorescence intensity, grouped by treatment condition and normalised relative to Vehicle (average of 1.5h and 2h timepoints). Datapoints and error bars indicate mean ± S.E.M. (D) Quantified time course data of proportion of cells exhibiting phagocytosis, grouped by treatment condition and normalised relative to Vehicle (average of 1.5h and 2h timepoints). Datapoints and error bars indicate mean ± S.E.M. (E) Grouped and individual data from last 3 timepoints of (C). ** < 0.01, *** < 0.005, **** < 0.0001, two-way ANOVA and post-hoc Tukey’s multiple comparison test. (F) Grouped and individual data from last 3 timepoints of (D). ** < 0.01, *** < 0.005, **** < 0.0001, two-way ANOVA and post-hoc Tukey’s multiple comparison test.

### EGCG dose-dependently inhibits microglia phagocytosis

We tested whether EGCG dose-dependently inhibited the microglial uptake of Aβ_42_. Microglia were treated with EGCG at either 0µM, 0.1µM, 1µM, or 10µM for 24h and then exposed to 500nM fluor 488-labelled Aβ_42_. Phagocytosis was monitored by fluorescence imaging as before (**Fig. 3A**). A significant effect of EGCG concentration was observed when monitoring the average 488-fluorescence intensity (mixed effects analysis treatment effect: F(3,20) = 3.85, P = 0.025, **Fig. 3B**) but not for the proportion of phagocytosing microglia (mixed effects analysis treatment effect: F(3,20) = 0.95, P = 0.44, **Fig. 3C**). Only 10µM EGCG at 1.5h and 2h showed a statistically significant difference from 0µM (Tukey’s multiple comparison 0µM vs 10µM: 1.5h P = 0.011, 2h P = 0.03) though there was a clear tendency in the data for a dose-response effect. The non-linear least squares method was used to fit a regression model to the normalised 488-fluorescence intensity at 2h (R^2^ = 0.51), showing a best-fit IC50 of 2.3µM for EGCG relative inhibition of microglia phagocytosis in this assay (**Fig. 3D**).

**Figure 3.**
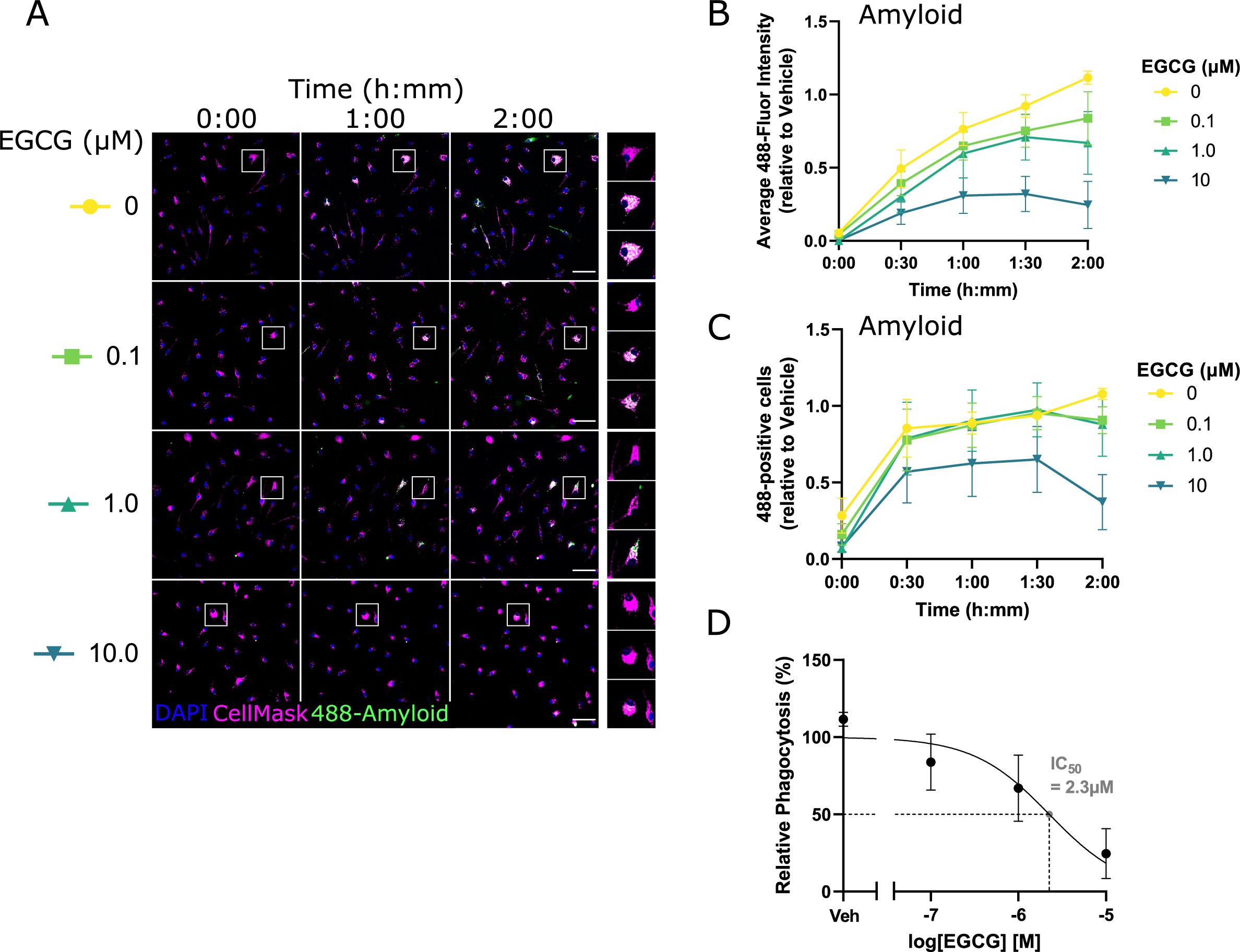
Dose-response screen for EGCG modulation of microglia phagocytosis. (A) Representative immunofluorescence images of primary microglia live-stained with a nuclear marker (DAPI; blue) and a cell membrane marker (CellMask; magenta), showing phagocytosis of HiLyte Fluor488 labelled Aβ_42_ peptide (green) at different timepoints. Individual rows represent individual EGCG concentrations. White boxes indicate inset images on the right panel. Scale bar = 50µm. (B) Quantified time course data of average cellular 488-fluorescence intensity, grouped by EGCG concentration and normalised relative to Vehicle (average of 1.5h and 2h timepoints). Datapoints and error bars indicate mean ± S.E.M. (C) Quantified time course data of proportion of cells exhibiting phagocytosis, grouped by EGCG concentration and normalised relative to Vehicle (average of 1.5h and 2h timepoints). Datapoints and error bars indicate mean ± S.E.M. (D) Concentration-response curve for EGCG inhibition of microglia phagocytosis.

### EGCG blocks microglia phagocytosis of Zymosan bioparticles

The inhibitory effect of EGCG on microglia phagocytosis of Aβ_42_ may derive from a direct effect of EGCG on microglia, and/or an effect of EGCG on Aβ_42_ peptide behaviour. The latter possibility is given credence by the reported metal chelating properties of EGCG [16], which could limit the aggregation, conformational transitions, and redox activity of Aβ_42_ [17,18] and thereby reduce its internalisation. To test this, we exposed EGCG-(10µM) or Vehicle-(0µM) treated microglia to fluorescent Zymosan bioparticles, which do not display amyloidogenic properties, and monitored their uptake over time (**Fig. 4A**). Zymosan bioparticles are derived from the yeast cell wall and are commonly used pathogens to induce sterile inflammatory responses and phagocytosis by host immune cells. Our data show that EGCG treatment strongly inhibited the time-dependent increase in microglia 488-fluorescence (mixed effects analysis EGCG effect: F(1,10) = 29.05, P=0.0003) with statistically significant differences observed from 1h onwards (Šídák’s multiple comparisons 0µM vs 10µM: 1h P = 0.011, 1.5h P = 0.002, 2h P = 0.0025, **Fig. 4B**). Similar, but less pronounced effects of EGCG were observed for the relative proportion of phagocytic microglia (mixed effects analysis EGCG effect: F(1,10) = 5.85, P=0.036) though none of the timepoints showed a significant treatment effect in post-hoc comparisons (**Fig. 4C**).

**Figure 4.**
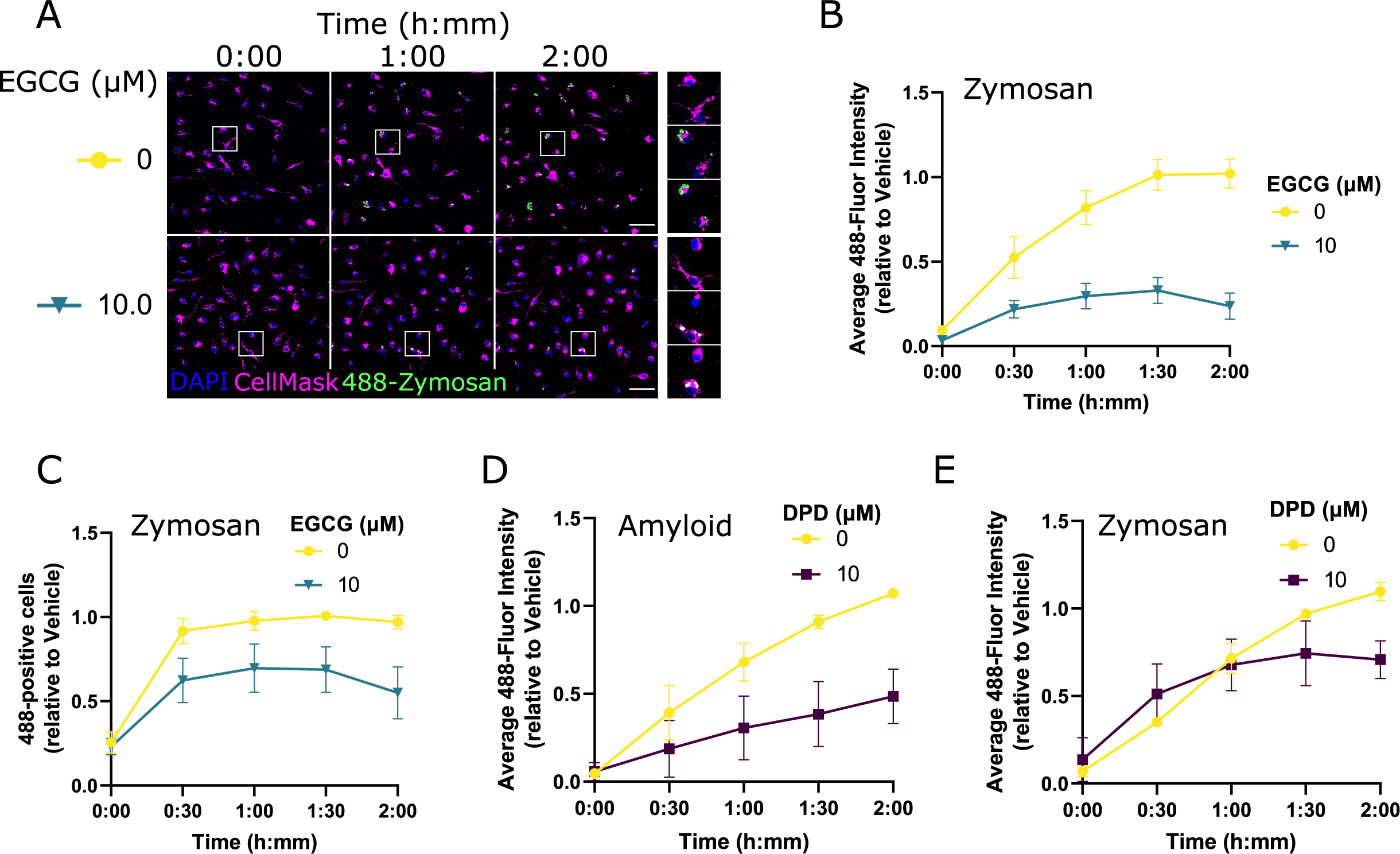
EGCG blocks microglia phagocytosis of Zymosan particles. (A) Representative immunofluorescence images of primary microglia live-stained with a nuclear marker (DAPI; blue) and a cell membrane marker (CellMask; magenta), showing phagocytosis of pHrodo Green Zymosan bioparticles (green) at different timepoints. Individual rows represent individual EGCG concentrations. White boxes indicate inset images on the right panel. Scale bar = 50µm. (B) Quantified time course data of average cellular 488-fluorescence intensity, grouped by EGCG concentration and normalised relative to Vehicle (average of 1.5h and 2h timepoints). Datapoints and error bars indicate mean ± S.E.M. (C) Quantified time course data of proportion of cells exhibiting phagocytosis, grouped by EGCG concentration and normalised relative to Vehicle (average of 1.5h and 2h timepoints). Datapoints and error bars indicate mean ± S.E.M. (D) Quantified time course data of average cellular 488-fluorescence intensity for 2,2’-Dipyridyl treated microglia internalising HiLyte Fluor488 labelled Aβ_42_ peptide, normalised relative to Vehicle (average of 1.5h and 2h timepoints). Datapoints and error bars indicate mean ± S.E.M. (C) Quantified time course data of average cellular 488-fluorescence intensity for 2,2’-Dipyridyl treated microglia internalising pHrodo Green Zymosan bioparticles, normalised relative to Vehicle (average of 1.5h and 2h timepoints). Datapoints and error bars indicate mean ± S.E.M.

These data indicate that EGCG affects the microglia phagocytosis of multiple pathogens and is not an effect specific to Aβ_42_. To validate this further, we compared the effects of a metal chelating compound, 2,2’-Dipyridyl (DPD), on microglia phagocytosis. DPD significantly reduced the uptake of Aβ_42_ in microglia, in a time dependent manner (mixed effects analysis DPD effect: F(1,4) = 6.8, P=0.059; mixed effects analysis DPD x time effect: F(4,15) = 4.8, P=0.011) though none of the timepoints showed a significant treatment effect in post-hoc comparisons (**Fig. 4D**). In contrast, no effect of DPD, was observed when monitoring microglia phagocytosis of Zymosan particles (mixed effects analysis DPD effect: F(1,17) = 2.2, P=0.16). The difference in effects of EGCG and DPD on Zymosan uptake by microglia therefore supports the idea that EGCG is modulating microglia phagocytosis independently from its metal-chelating properties (**Fig. 4E**).

### EGCG modulates inflammatory cytokine release from microglia

The phagocytosis assay data indicated that EGCG is likely acting directly on microglia to modulate cell function. Given that microglia are well known mediators of inflammatory signalling in the brain [19], we tested whether EGCG affects the release of inflammatory cytokines from microglia in culture.

Microglia conditioned media (MCM) following 24h exposure to 1µM EGCG was examined using the Mouse XL Proteome Profiler Array, which semi-quantitatively measures the abundance of 111 different cytokines released into the media. For comparisons, MCM was also examined from microglia treated with Vehicle, or with the established pro-inflammatory and anti-inflammatory stimuli IL-3 and IL-4, respectively [20,21]. Quantification of dot blot intensity and normalisation of each cytokine relative to Vehicle, showed that EGCG treatment of microglia reduced the levels of nearly all cytokines in the MCM (**Fig. 5A, B**). Hierarchical clustering of cytokines by their changes to expression levels revealed subsets of proteins, including IL-17a, CCL20, IGFBP-5, Complement Factor D and IL-22, that were most strongly suppressed by EGCG treatment (**Fig. 5C, D**). Protein-protein interactions among the EGCG-suppressed proteins were extracted from the STRING database (**Fig. 5E**) and queried through g:Profiler (GO:BP with < 300 terms) [22], which highlighted immune cell migration and chemotaxis (e.g., leukocyte chemotaxis, Padj = 1.963×10^-27^ among several immune processes strongly associated with these protein networks. Together, these data suggest that EGCG suppresses immune cell signalling cues used by microglia to elicit inflammatory responses.

**Figure 5.**
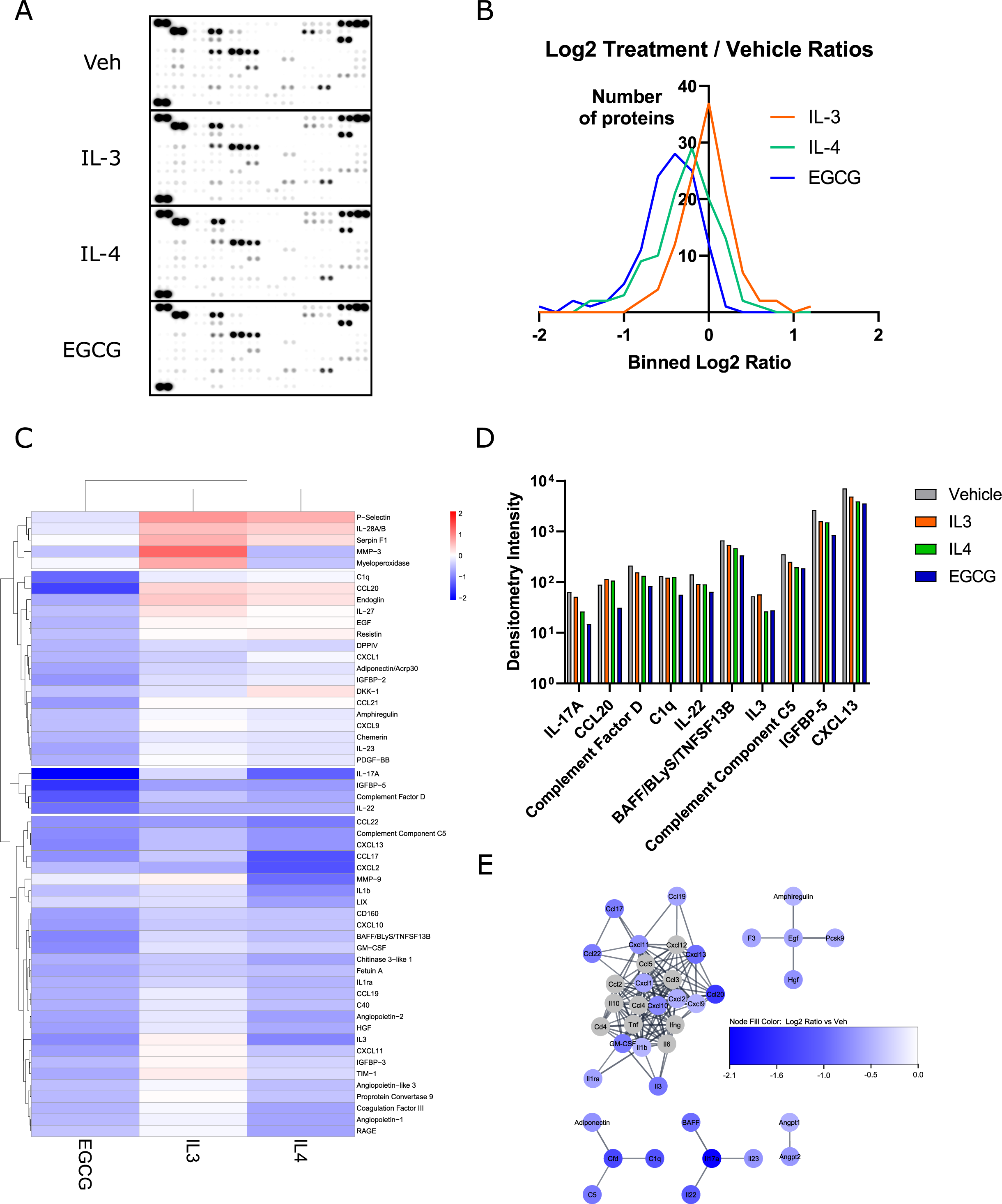
EGCG reduces inflammatory cytokine release from microglia. (A) Dot blots of 111 different cytokines (spotted in duplicate) of the Proteome Profiler Mouse XL Cytokine Array exposed to conditioned media from microglia treated with EGCG (1µM) or IL-3 (20ng/ml) or IL-4 (20ng/ml) for 24h. (B) Log2 ratios of densitometric readings for all 111 cytokines from (A) for each treatment relative to Vehicle. (C) Heatmap and hierarchical clustering of cytokines showing an absolute log2 Treatment/Vehicle ratio > 0.5 in any of the treatment conditions. Colour indicates log2 Treatment/Vehicle ratio. (D) Densitometry values of the top 10 cytokines modified by EGCG relative to Vehicle. (E) STRING network analysis of functional protein associations between the top EGCG modified cytokines. Node colour indicates Log2 EGCG/Vehicle densitometry ratio for each cytokine (grey nodes are functionally associated proteins that were not changed/tested in the Profiler Array). Edges represent predicted protein associations with stringdb confidence scores > 0.8 (mapped to line width).

### EGCG modifies the microglia transcriptome

RNA sequencing was used to examine the transcriptome of microglia treated for 24h with 1µM EGCG (N=2). 136 differentially expressed genes were identified relative to vehicle-treated microglia, with a non-adjusted p-value <0.05, including 81 genes that were up-regulated and 55 genes that were down-regulated (**Fig. 6A**). 6 of these genes were differentially expressed with an adjusted p-value < 0.1, including 4 genes that were up-regulated by EGCG (*Rpl17, Cd36, Plau, MgII*) and 2 genes that were down-regulated (*Serpinb2, Mmp3*) (**Fig. 6A**).

**Figure 6.**
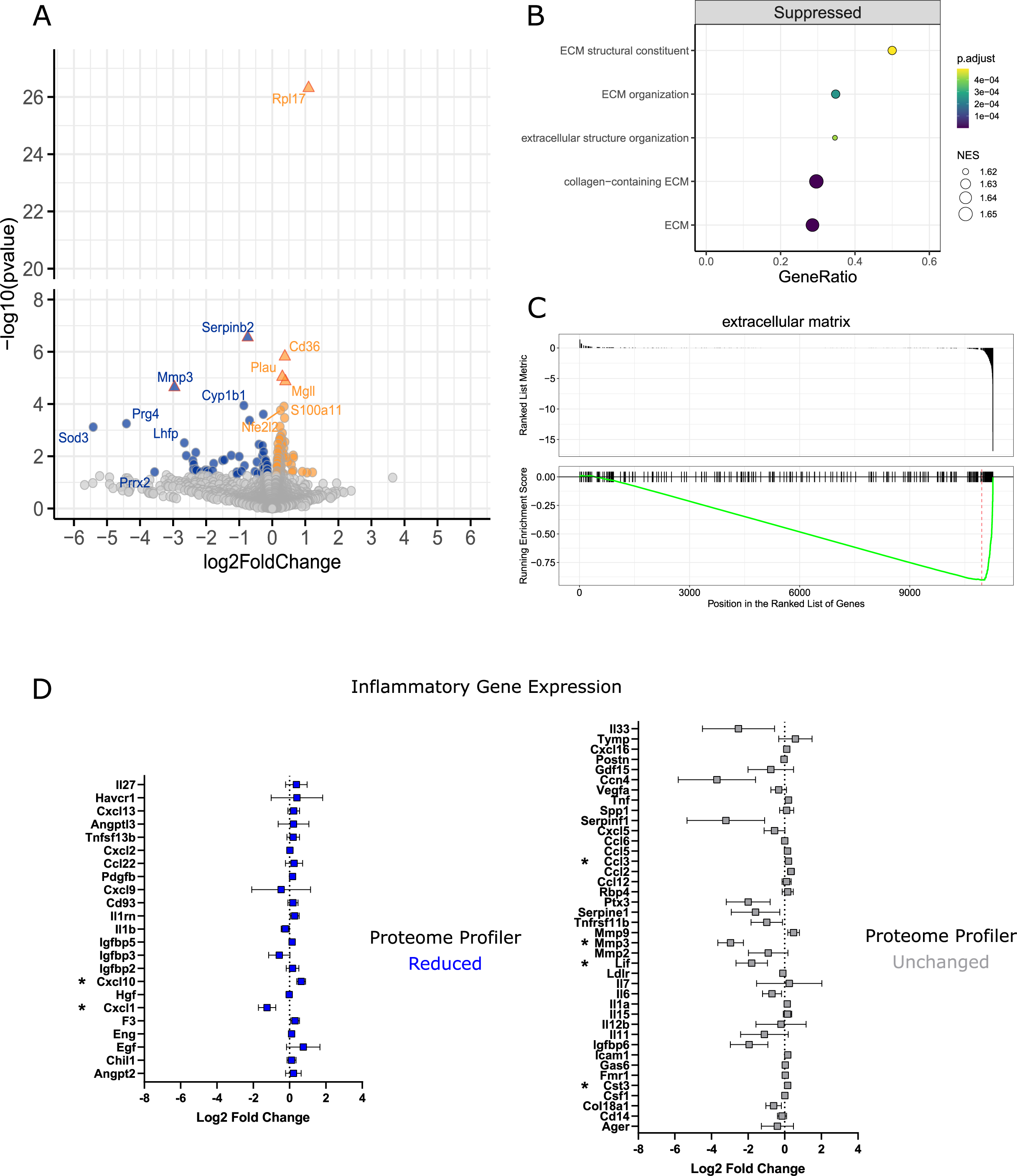
EGCG modifies the microglia transcriptome. (A) Volcano plot showing differentially expressed genes (DEGs) in EGCG treated microglia compared to Vehicle. DEGs with a nominal p-value < 0.05 are coloured in blue (downregulated) or orange (upregulated). DEGs with a BH-adjusted p-value < 0.1 are shown as triangle symbols with red outlines. (B) Gene sets significantly suppressed in EGCG treated microglia with Normalised Enrichment Scores (NES) and Gene Ratios (proportion of DEGs within gene set size). ECM = extracellular matrix. (C) GSEA plot of ECM genes according to their ranked position within the list of EGCG modified genes. (D) EGCG-induced changes to transcription of inflammatory genes that show reduced (log2 densitometry ratio < - 0.5; left panel) or unchanged (log2 densitometry ratio > - 0.5; right panel) protein release in the Proteome Profiler Array. Gene expression changes with nominal p-values < 0.05 are marked by an asterisk.

To provide further biological context to the EGCG induced transcriptome changes, we applied Gene Set Enrichment Analysis (GSEA) to the ranked list of EGCG-modified genes. This approach highlighted that genes involved in the composition and regulation of the microglia extracellular matrix (ECM) were significantly downregulated by EGCG (**Fig. 6B, C**).

Comparing gene expression level changes for inflammatory proteins analysed in MCM (**Fig. 4**) revealed little correlation between gene and protein-level changes. Only 2 genes with reduced MCM protein levels were significantly altered in the microglia transcriptome by EGCG; *Cxcl1* (down-regulated) and *Cxcl10* (up-regulated) (**Fig. 6D**). Conversely, several genes with unchanged MCM protein levels were significantly changed in the microglia transcriptome by EGCG; *Lif, Mmp3* (down-regulated) and *Ccl3, Cst3* (up-regulated). These data indicate that the strong modulatory effects of EGCG on inflammatory cytokine release are not likely to be through a direct transcriptional mechanism.

## Discussion

Microglia are crucial players in the pathogenesis of late onset AD, with evidence for both deleterious and beneficial effects [23]. The proliferation and activation of microglia close to Aβ plaques is consistent with the prevailing view of a functional removal of toxic protein aggregates, although engulfment of stressed but still viable synapses is also likely [24]. Plaques fail to form in the parenchymal space of microglia knockout mice so microglia could contribute to the development and spread of Aβ pathology [3]. Therefore, understanding how AD intervention strategies impact on microglial function is crucial when assessing overall therapeutic potential. Due to growing interest in the beneficial actions of dietary flavanols on brain health, combined with the progression of EGCG into clinical trials for AD, we embarked on structure function studies on a small series of related flavan-3-ols to define their impact on microglial responsiveness. Because our primary interests are in risk reduction strategies for late onset AD, we chose not to use APP or PS1 expression models, or microglia derived from human iPSc carrying mutations instead preferring well-established, and fully characterised primary rodent microglia. These mouse-derived cells expressed C11b and Iba1 and exhibited key functional characteristics such as NFκB translocation, NO release, and Aβ phagocytosis as expected of microglia in the CNS. Applying flavanol-3-ols to these cells and assessing responsiveness indicated that the extent and pattern of hydroxylation and the presence of the galloyl moiety were strong determinants of the microglia modifying properties of flavan-3-ols. EGCG was the most effective flavan-3-ol inhibiting phagocytosis of Aβ, demonstrating broadly anti-inflammatory activity, reducing cytokine release and downregulating transcription particularly of components of the microglia extracellular matrix.

EGCG showed a clear time- and concentration-dependent inhibitory effect on the phagocytosis of Aβ. This could have resulted from direct modulatory actions on microglial reactivity or, could have been secondary to molecular interactions between EGCG and Aβ. Aβ aggregation, redox activity and toxicity are closely associated with the binding of iron and other metal ions and EGCG is a well characterised metal chelator, antioxidant [16] and disrupter of Aβ aggregation [11] so we considered this as a potential mechanism of action. Indeed, the bidentate chelating compound DPD, which blocks Aβ toxicity in neurons [25] also inhibited Aβ phagocytosis in our microglial model supporting the role of iron chelation as an underlying mechanism. However, EGCG also strongly inhibited the phagocytosis of zymosan which was not influenced by metal chelation. This suggests that EGCG is acting to influence microglial phagocytosis through at least another mechanism, potentially through modulation of receptor signalling pathways [26]. For example, flavan-3-ols are well characterised inhibitors of key protein kinase pathways lying downstream of TAM receptors [27]. Indeed, the TREM2-DAP12 complex is strongly implicated in microglial responsiveness in AD, co-ordinating signaling via Syk to PI3-kinase, Akt and ERK [28] all of which are potential targets for flavan-3-ols. Significantly, TAM receptor-driven microglial phagocytosis does not inhibit, but promotes, Aβ plaque development [2] so flavan-3-ol inhibition of Aβ phagocytosis through such a mechanism could block dense-core plaque formation.

To relate flavan-3-ol inhibition of phagocytosis to the overall inflammatory status, microglia were treated with EGCG and assessed for cytokine release in comparison with IL-3 (pro-inflammatory) and IL-4 (anti-inflammatory) using a proteome profiler. Although an anti-inflammatory phenotype was not unexpected for EGCG [29] the extent and breadth of inhibition of cytokine release with EGCG was very notable particularly in comparison to that seen with IL-4. Clustering of affected cytokines revealed subsets including IL-17a, CCL20, IGFBP-5, Complement Factor D, and IL-22, that were all very strongly suppressed by EGCG treatment. Further analysis highlighted immune cell migration and chemotaxis among several immune processes most strongly associated with these protein networks. Together, these data suggest that EGCG suppresses immune cell signalling cues used by microglia to elicit inflammatory responses with the possibility therefore of modulating migration to Aβ plaques.

To gain further functional insight into the flavanol-3-ol response, RNA sequencing was undertaken to define the EGCG microglial transcriptome. There was surprisingly little correlation between the RNA sequencing and proteome profiler data sets and from this we conclude that most changes in the EGCG-evoked cytokine secretome did not result from direct transcriptional responses, although some changes might have been missed due to the temporal nature of gene expression. Genes upregulated by EGCG treatment included the ribosomal protein Rpl17, the urokinase-plasminogen activator gene Plau, which has been implicated in AD risk [30] and most notably the cell surface scavenger receptor CD36 which binds Aβ fibrils as part of a phagocytic response [31]. With regards to the identified down regulated genes, microglial matrix metalloproteinases (MMPs) such as MMP3 are important mediators of neuroinflammation and synaptic reorganisation in plasticity and neurodegeneration [32]. Pharmacological inhibition of MMP3 has previously been postulated as an intervention for inflammatory diseases of the CNS [33], so EGCG might have therapeutic potential in this respect. SerpinB2, also known as type 2 plasminogen activator inhibitor (PAI-2) is upregulated in activated microglia in AD [34] and may play a role in migration and matrix breakdown. More recent data proposes a role for SerpinB2 in protein misfolding and proteostasis [35,36] so the functional consequence of EGCG mediated downregulation of SerpinB2 is difficult to predict.

Increasing levels of recognition of the importance of microglia in AD and other forms of neurodegeneration highlights the potential need for modulators of phagocytosis and brain inflammation to slow onset and progression. Bioavailable flavan-3-ols may hold some promise but the extent of hydroxylation and a galloyl moiety appear strong determinants of function which will need careful consideration with respect to ADME and future drug design based around flavanol scaffolds.

## Author contributions

PR and RJW conceived and designed the study. PR, KLH and JS contributed to material preparation and provided technical advice. PR conducted the experiments and undertook data collection and analysis. PR and RJW wrote the manuscript. All authors read and approved the final manuscript.

## Acknowledgements

PR and RJW were supported by an Alzheimer’s Society Project Grant (AS-PG-18b-009). PR was further supported by an Alzheimer’s Research UK Pump Priming Grant and a GW4 Generator Fund Grant (GW4-GF2-004) and RJW. by an Alzheimer’s Research UK Equipment Grant (ARUK-EG2018A-008). KLH was supported by grant MR/N0137941/1 for the GW4 Biomed MRC DTP, awarded to the Universities of Bath, Bristol, Cardiff, and Exeter from the Medical Research Council (MRC)/UKRI.

